# Rapid and sensitive on-site genetic diagnostics of pest fruit flies using CRISPR-Cas12a

**DOI:** 10.1101/2022.06.22.497159

**Authors:** Dan Mark Alon, Tamir Partosh, David Burstein, Gur Pines

**Affiliations:** Department of Entomology, Agricultural Research Organization - the Volcani Center, 68 HaMaccabim Rd, Rishon LeZion 7505101, Israel; The Shmunis School of Molecular Cell Biology & Biotechnology, Faculty of Life Science, Tel Aviv University, Tel Aviv 69978, Israel

**Keywords:** pest-control, CRISPR-Cas12a, RPA, genetic detection, *Bactrocera zonata*, *Ceratitis capitata*

## Abstract

*Bactrocera zonata*, a major fruit pest species, is gradually spreading west from its native habitat in East Asia. In recent years it has become a major threat to the Mediterranean area, with the potential of invading Europe, the Americas, and Australia. To prevent its spreading, monitoring efforts in plantation sites and border controls are carried out. Despite these efforts, and due to morphological similarities between *B. zonata* and other pests in relevant developmental stages, the monitoring process is challenging, time-consuming, and requires external assistance from professional labs. CRISPR-Cas12a genetic diagnostics has been rapidly developing in recent years and provides an efficient tool for the genetic identification of pathogens, viruses, and other genetic targets. Here we design a CRISPR-Cas12a detection assay that differentially detects two major pest species, *B. zonata* and *Ceratitis capitata.* Our easy-to-use and affordable assay employs a simple DNA extraction technique together with isothermal amplification, and Cas12a-based detection. We demonstrate the specificity and high sensitivity of this method, and its relevance for on-site applications. This method is highly modular, and the presented target design method can be applied to a wide array of pests.

**Key Massage:**

- Distinguishing different pest fruit flies on-site is crucial for prevention of global spreading but can be difficult
- We present a genetic identification assay for rapid, on-site detection of pest using CRISPR-Cas12a
- The method is affordable, quick and easy-to-use, and can be applied in border controls or on-site
- The design process can be easily tailored for any pest, and can greatly benefit developing countries

## Introduction

In recent years, the peach fruit fly, *Bactrocera zonata* (Saunders, Diptera: Tephritidae) has become a major invasive species in Africa and the Arab peninsula. Globally, it is responsible for annual losses of hundreds of millions of USD (EPPO 2005; Stonehouse, Mumford, and Mustafa 1998). *Bactrocera zonata* is an aggressive, highly adaptable invasive species with a broad range of host plants, covering more than 50 commercial and wild plant species, mainly fleshy fruits and vegetables (EPPO 2010). This fly is native to East and South-East Asia and hence well accustomed to tropical and subtropical climates. Nonetheless, it was shown to establish in colder climates reaching freezing-point, enabling its proliferation in the Mediterranean climate (Delrio and Cocco 2012). Combined with global warming, this makes *B. zonata* a serious threat to West Asia, several European countries as well as parts of Australia and the Americas (Zingore et al. 2020; Rosace et al. 2019).

*B. zonata* is listed as an A1 pest in the European and Mediterranean Plant Protection Organization (EPPO) and is regulated by many EPPO member countries (EPPO 2010). The ability to accurately detect different insect pests is a vital first step in the combat against the global spreading of invasive pest species. This effort typically takes place in border controls and involves the investigation of eggs and larvae found in fruits and vegetables. Morphological identification in these early life stages is a major tool used to distinguish between different species, but can be problematic due to high similarities between species, such as in Israel, with *B. zonata* and the much more abundant species *Ceratitis capitata* (Figure 1a). These similarities require specimens to be reared to adulthood for identification. Such practices can be hazardous as not all countries possess adequate quarantines. Moreover, rearing eggs is timely and often unfruitful (Armstrong et al. 1997). Aside from the risks involved in rearing, during this process, the entire cargo is halted and could go to waste, leading to considerable losses both to farmers and customers.

**Figure 1.**
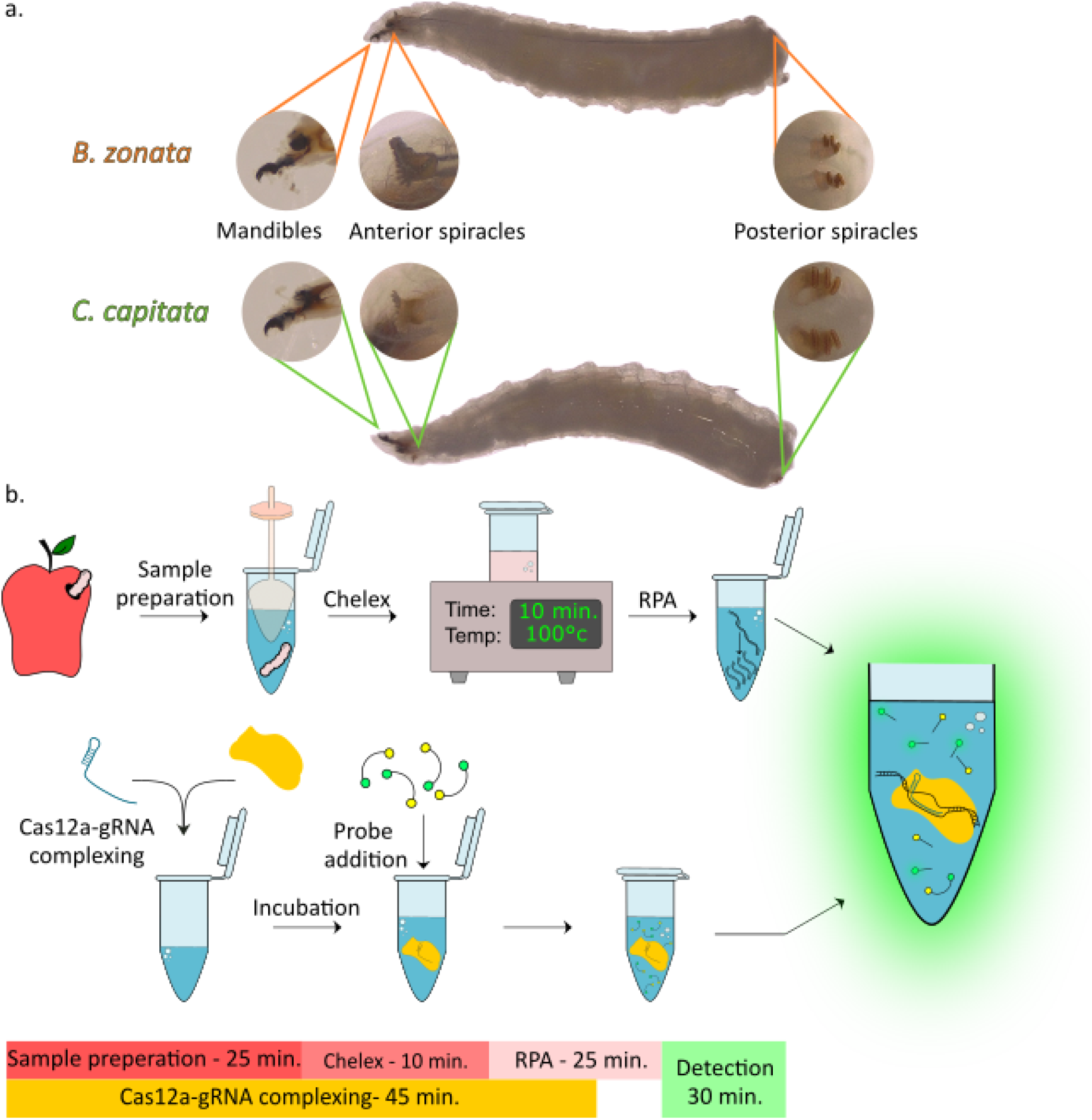
Scheme of proposed assay (a) Comparison of the organs used for larval stage morphological identification of *B. zonata* (top) and *C. capitata* (bottom). (b) A scheme of the detection assay. After sample collection from the fruit, larvae are homogenized, Chelex 100 resin was added to the homogenate and boiled for 10 min. Next, 2 μL of the solution is amplified by RPA. In parallel, a Cas12a-gRNA complex is assembled (or thawed), a reporter and the amplicons are added to the solution and incubated for 30 min. Finally, fluorescence is measured to test for the presence of the target DNA.

Extensive efforts have been made to facilitate faster, more accurate diagnostic techniques. Molecular methods for the identification of different *Bactrocera* species have been developed to aid in the battle against the spread of these harmful pests. Species-specific markers based on the mitochondrial cytochrome oxidase COII gene (Asokan et al. 2011), restriction fragment length polymorphism (RFLP) detected in a polymerase chain reaction (PCR), amplified ribosomal DNA (rDNA)(Armstrong et al. 1997), and high-resolution melt (HRM) real-time PCR assays (Dhami and Kumarasinghe 2014) have been developed and are currently being used to distinguish between different pests. These methods have considerably reduced the quarantine time but still depend on the availability of lab space, technicians, expensive instruments, and reagents. Such methods are usually performed off-site, requiring specimen transportation to specialized facilities, thus increasing the handling time and raising the probability of spreading events.

Recently, CRISPR-Cas-based diagnostic approaches have provided a new set of accurate detection tools at the molecular level. Specifically, a type V CRISPR-Cas enzyme, *Lachnospiraceae bacterium* ND2006 Cas12a (LbCas12a), presents great promise in the field of *in-situ* molecular detection of viruses, pathogens or any DNA sequence of interest (Alon et al. 2021; Wu et al. 2021; T. Zhang et al. 2021; Broughton, Deng, Yu, Fasching,Singh, et al. 2020). LbCas12a is an RNA-guided enzyme capable of recognizing double-stranded DNA (dsDNA) targets and performing a protospacer adjacent motif (PAM)-distal dsDNA break with staggered ends (Zetsche et al. 2015). Upon detection of a dsDNA target, the enzyme is activated, and in addition to its target-specific cleavage activity, it initiates indiscriminate single-stranded DNA (ssDNA) cleavage (Abudayyeh et al. 2016, 2;East-Seletsky et al. 2016, 2). This attribute was found to be very effective for DNA detection (Chen et al. 2018; Kellner et al. 2019). LbCas12a-based DNA detection requires a modified ssDNA reporter, usually containing a fluorophore and a quencher to be added to the detection reaction. Once activated, the LbCas12 cleaves the ssDNA reporter, releasing the fluorophore from the quencher, leading to an easily-detectable fluorescence signal (Figure 1b). The detection reaction only requires a short incubation at 37°C, and its components can be lyophilized to increase stability during storage. With minor variations, detection can be held using lateral flow strips, thus eliminating the need for a fluorescence-reading device (Broughton, Deng, Yu, Fasching, Singh, et al. 2020; Curti et al. 2021).

Most DNA detection methods require a DNA amplification step, which might be vital if DNA material is scarce. Different methods are used to amplify the target sequence prior to detection. PCR amplification is the gold-standard method for DNA amplification and has been demonstrated to work well with the Cas12a detection system (Alon et al. 2021; Liu et al. 2022). Two leading isothermal methods are commonly used for in-situ applications: loop-mediated isothermal amplification (LAMP) and recombinase polymerase amplification (RPA). LAMP is based on a strand-displacing polymerase with 4-6 primers, and enables the amplification of short DNA segments (80-250bp) in ~30 minutes, at 60-65°C (Notomi et al.2000). RPA is based on an ssDNA binding protein (SSB), a recombinase that leads the SSB-primer complex to its destination, and a strand-displacing polymerase that initiates an isothermal amplification reaction. RPA requires only two primers, performs better than PCR with longer-sized primers, and offers fewer constraints on primer design, allowing higher GC content (Piepenburg et al. 2006). RPA reactions are 10-20 min. long, are performed at 37-42°C and offer high sensitivity even with minuscule amounts of starting material (Aman et al. 2020). Amplification of a pool of several specimens is of great benefit since co-infestation of fruit by different Tephritidae and Drosophilidae families occur regularly (Deus et al. 2016;Zida et al. 2020; Moquet et al. 2021). Therefore, an amplification method that allows the pooling of different specimens as well as multiplexed detection of the different targets in the pool can prove beneficial for simple *in-situ* detection of pests.

In this work, we present a rapid and simple method to accurately differentiate between two highly hazardous pest species: *B. zonata* and *C. capitata.* We have designed an on-site assay that utilizes a simple DNA purification technique together with RPA amplification and LbCas12a detection. Our assay requires only a hot plate, a bench-top centrifuge, and a hand-held fluorometer, yet it can distinguish between the two pests with high confidence in a little over an hour. We believe this assay can aid in the battle against invasive species global spreading, avoiding quarantines and losses for farmers and consumers. Our assay is modular and can be easily expanded to other species, making it relevant for pests of concern worldwide. Notably, the affordability and simplicity of this method make it uniquely suited for pest control in developing countries.

## Results

### Protocol outline

Our protocol consists of three main steps: (a) Sample collection and DNA extraction using the Chelex 100 resin, (b) an RPA amplification of specific targets, and (c) Cas12a detection. As illustrated in Figure 1b, the entire protocol takes roughly one hour and a half to complete, most of which consists of incubation rather than hands-on time. The protocol requires only a benchtop centrifuge and a hot-plate. In this work, the readout was obtained using a plate reader for throughput purposes, but alternative methods such as lateral-flow-based assays or handheld field fluorometers are also applicable (Broughton, Deng, Yu, Fasching,Servellita, et al. 2020; Kurosaki et al. 2016).

### Pest-specific identification

Specific target selection for *B. zonata* was performed through pairwise sequence alignment of the mitochondrial genome sequences of *B. zonata* and *C. capitata* (Spanos et al. 2000;Choudhary et al. 2015; Katoh, Rozewicki, and Yamada 2019). We used *C. capitata* as a reference genome for a non-target fly since its larvae are morphologically very similar and cannot be easily differentiated (Figure 1a). We sought variable regions within the mitochondrial genomes of both insects, and used one such region (VR1 - Figure 2a, Table 1) to design primers and guide RNAs (gRNAs) specific to *B. zonata*.

**Figure 2.**
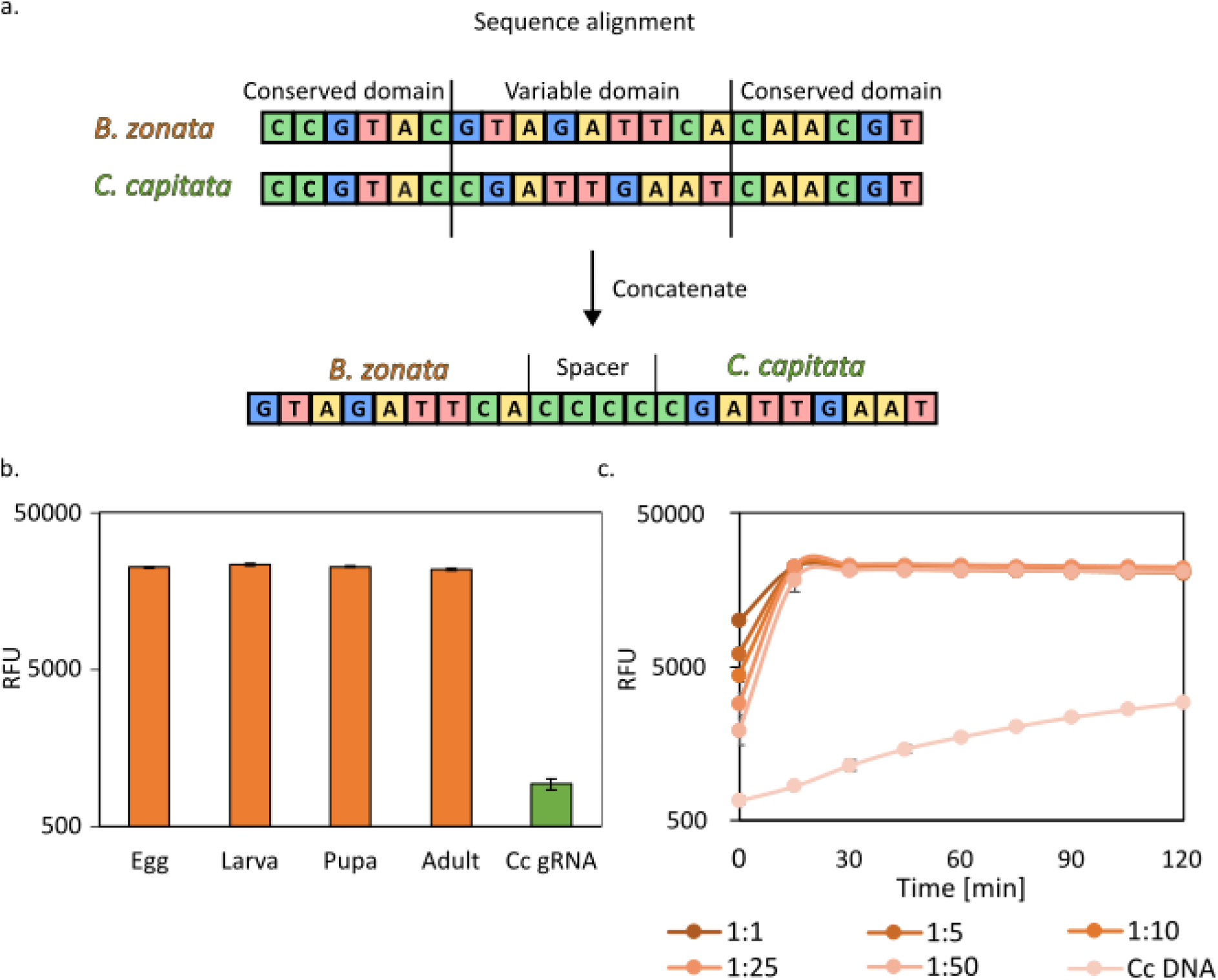
Pest-specific identification. (a) A scheme of the gRNA design process. Mitochondrial sequences of relevant pests were aligned and variable regions were identified and concatenated for gRNA design using CRISPOR. The uniqueness of the gRNA sequences was then validated and tested experimentally (Figure S1). (b) Cas12a detection of *B. zonata* in samples of different developmental stages (orange). DNA was obtained from fresh samples using Chelex 100 (Methods). Amplification was performed using RPA with specific primers for *B. zonata* and Bz1 gRNA (Table S1) was used for detection (Table S1). As a negative control, *C. capitata* larvae were used with Bz1 gRNA (green). (c) Sensitivity of Cas12a-Bz gRNA detection in varying ratios of pooled larvae. Samples containing a single larva of *B. zonata* with increasing amounts of *C. capitata* larvae were prepared. DNA was then extracted and diagnosed as in (b), using *B. zonata* specific primers and Cas12a-Bz gRNA complexes. As a negative control, *C. capitata* DNA was used. All experiments were performed with three biological repeats and three technical repeats.

First, we designed several gRNAs using the CRISPOR online platform (Concordet and Haeussler 2018). To ensure the gRNAs are specific to our target *B. zonata* only, we concatenated both variable regions from *B. zonata* and *C. capitata* and used the merged sequence as a template for CRISPOR. We have also added a spacer between the sequences in order to avoid the selection of gRNAs derived from the stitch between the two variable regions. We selected the best-scored hits and double-checked the sequences to verify no cross-reactivity exists. Based on the gRNAs position within VR1, we designed specific primers for PCR and RPA amplification (Table S1). Next, we used a commercial DNA extraction protocol (see Methods section) on *B. zonata* and *C. capitata* adult flies, amplified our target with PCR, and tested three different gRNAs (Table 1 and Figure S1). Only one of the gRNA (Bz1) displayed a specific signal in the presence of *B. zonata’s* genome, and it was used for the follow-up analyses.

Next, we tested the full protocol on samples of both fly species: DNA was extracted from the samples using Chelex 100 resin. Chelex is a chelating resin with a high affinity for polyvalent ion metals, enabling simple and cheap DNA extraction. It has been shown to facilitate DNA extraction from cells at 100°C by chelating metal ions that act as DNase catalysts, thus preventing DNA degradation (Walsh, Metzger, and Higuchi 1991). With slight modification to the protocol, Chelex 100 can be used to efficiently extract DNA from insects with as little as a single egg for starting material (Musapa et al. 2013).

Following extraction, DNA was isothermally amplified using RPA and tested with Cas12a-gRNA complexes. We began by examining the ability to detect a positive and specific signal from all developmental stages of *B. zonata* (Figure 2b). The Chelex 100 DNA extraction method proved to be highly efficient throughout all developmental stages of the flies, including single eggs, as demonstrated previously. The RPA efficiently amplified the Chelex-originated DNA material in only 20 minutes at 37°C. The complexes consisting of the Cas12 and the gRNAs we designed properly detected their targets and were inactive on the non-targeted sequences.

Next, we tested the sensitivity of our assay. We combined larvae from both *B. zonata* and *C. capitata* in ratios ranging from a 1:1 ratio and up to 1:50 larvae, respectively (Figure 2c). A robust signal was observed in all ratios, and no signal was observed in the absence of *B. zonata* DNA. These results suggest that pooling larvae from several fruits is feasible without loss of specificity, allowing to cut down the required time, costs, and labor. We further tested for cross-reactivity between *B. zonata* and *Drosophila melanogaster* as a representative fermentation fruit fly. This test is relevant to real-life scenarios since opportunistic species such as *D. melanogaster* may lay eggs in damaged fruits after picking and shipment, leading to larvae in the pooled sample tube. We used the same ratios (1:1 up to 1:50), and no cross-reactivity was detected (Figure S2).

### Universal amplification assay

After validating the method using species-specific primers, we expanded the assay to include *C. capitata* identification using a universal RPA amplification reaction. By designing universal primers that can amplify variable regions in different fly species, we were able to produce more relevant data from a single pooled tube, while reducing the hands-on time and simplifying the protocol. We designed universal primers that match both *B. zonata* and *C. capitata* using the VR1 flanking regions (Figure 2a). Next, we picked the highest-ranking gRNAs for *C. capitata* from the CRISPOR predictions, and after sequence verification of all gRNA hits, we decided to proceed with only one gRNA (Cc2), as other hits were not specific to *C. capitata.*

The Chelex-extracted DNA from the samples described above was amplified, this time using the universal primers we designed and tested for the new gRNAs. First, we tested the ability to detect the targeted organisms in all developmental stages, as described above (Figure 3a). These results established that the universal primers are indeed capable of amplifying DNA from both species. Next, the sensitivity of the universal detection assay was examined, and again, we observed positive detection for both species with high confidence even at very low ratios of 1:50 (Figure 3b,c). We also cross-tested the gRNAs for the opposing RPA amplification products, and no cross-reactivity was detected. We did observe, however, a slight drop in the fluorescence signal in both reactions in the higher ratios (1:25 and 1:50) compared to the specific primers presented in Figure 2c. It is possible the lower signal is related to the higher amount of amplified DNA, as both insect species were amplified during RPA universal amplification.

**Figure 3.**
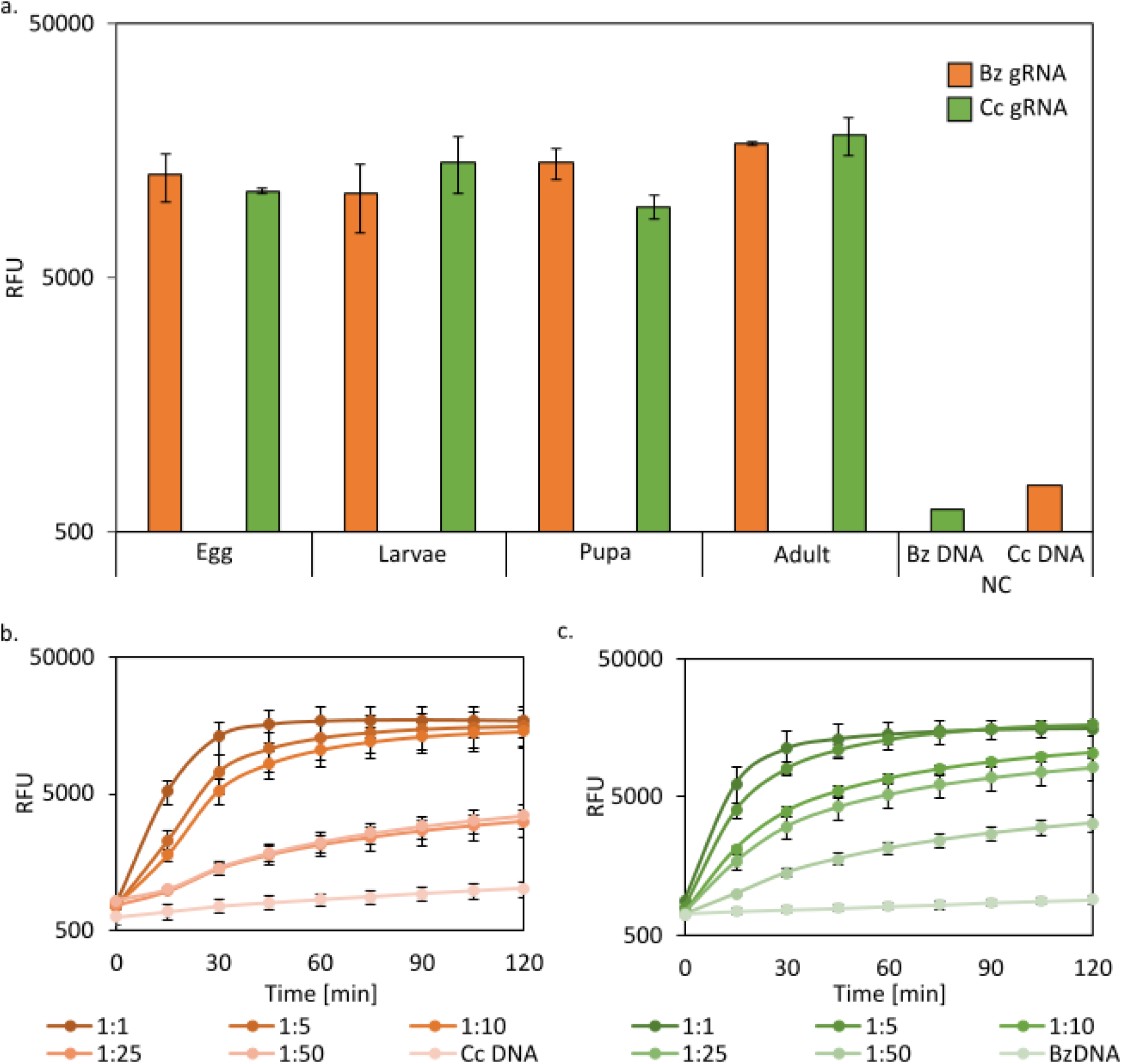
Universal amplification assay. (a) Cas12a detection of *B. zonata* (orange) and *C. capitata* (green) from different developmental stages. DNA was obtained using Chelex 100 from fresh samples (Methods). Amplification was performed using RPA with universal primers (Table S1) and detection was achieved using Bz1 gRNA for *B. zonata* and Cc2 for *C. capitata* (Table S1). As a negative control (NC), *C. capitata* and *B. zonata* larvae were used with non-corresponding gRNAs. (b) Detection sensitivity of Cas12a-Bz1 gRNA in varying ratios of pooled larvae. Samples containing a single larva of *B. zonata* with increasing amounts of *C. capitata* larvae were prepared. DNA was then extracted using Chelex 100, RPA amplified (using VR for\rev primers, Table S1) and diagnosed using Cas12a-Bz1 gRNA complexes. As a negative control, *C. capitata* DNA was used. (c) Detection sensitivity of Cas12a-Cc2 gRNA for *C. capitata* in varying ratios of pooled larvae. Samples containing a single larva of *C. capitata* with increasing amounts of *B. zonata* larvae were prepared. DNA was extracted and amplified as in (b), and diagnosed using Cas12a-Cc2 gRNA complexes. As a negative control, *B. zonata* DNA was used. All experiments were performed with three biological repeats and three technical repeats.

Lastly, we were also interested in assessing the stability of the detection and RPA reactions in storage. This aspect is important as long storage enables the preparation of bulk amounts of detection reactions in advance, reducing the amount of processing time per sample. We initially tested several conditions including room temperature, 4°C, and −20°C for a period of over 160 hours (Figure 4). Room temperature samples lost activity after 24 hours, and it appeared that evaporation took place, which might account for the loss of activity. Samples kept at 4°C lost activity between 48 and 168 hours and samples kept at −20°C showed no significant loss of activity within the period tested.

**Figure 4.**
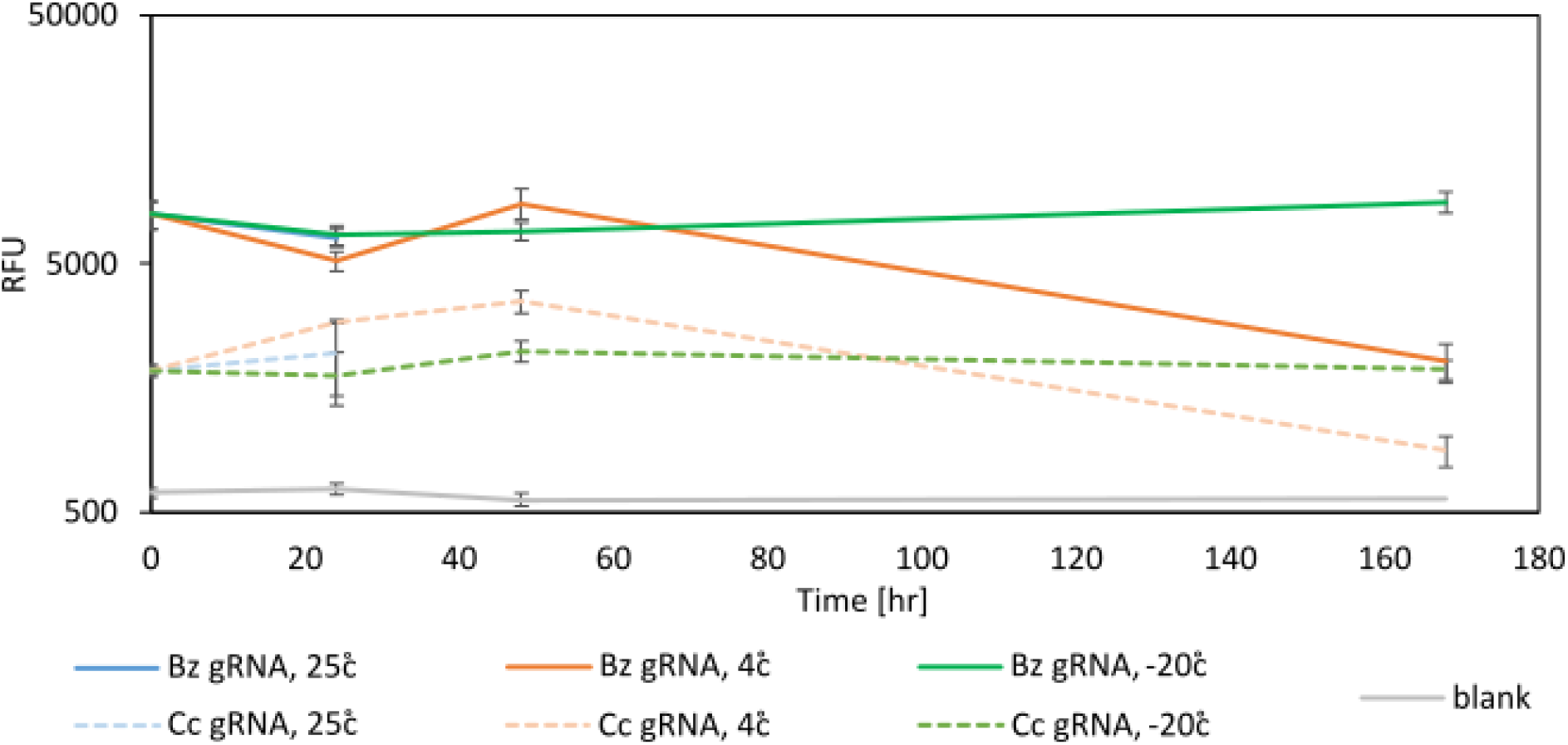
Stability of Cas12a complexes in different conditions. Samples were tested at time 0, after 24hr, 48hr and 167hr at room temperature, 4°C, and −20°C. *B. zonata* larvae RPA amplified samples were detected using Cas12-Bz1 gRNA complexes, and the same samples were detected with Cas12-Cc2 gRNA complexes (Table S1) as a negative control. Blank (gray) are detection reactions without DNA

## Discussion

In this work, we demonstrate a fast, reliable, straightforward, and affordable genetic detection assay for pest species, enabling accurate point-of-care pest control. Our assay requires only a bench-top centrifuge, a hot-plate, and a hand-held fluorometer, and can further be adapted for paper-based detection (Broughton, Deng, Yu, Fasching, Servellita, et al. 2020). The current price estimate per reaction is about 1.6 USD, and reagent prices may considerably decrease when purchased in bulk (Table S2).

We have demonstrated that the assay can be tailored to detect a specific fruit fly species or used for several species with the potential for universal amplification, including species that are highly similar, both morphologically and genetically. Universal reactions reduce hands-on time and enable the pooling of specimens in a single reaction. The Chelex method for DNA extraction has proven to be efficient in all developmental stages of the target flies and together with the robust RPA method, they allow the detection of even minute amounts of sample. The ability to distinguish a single target larva from a pooled tube containing up to 50X non-targets makes this assay meet and even surpass current gold-standard methods while also dramatically reducing labor and cost (Koohkanzade et al. 2018).

There are several limitations to our current setup. First, RPA reagents are currently sold by a single company creating a potential bottleneck in reagent supply. Second, as in all genetic-based methods, genomic data of target pests is crucial for gRNA design and target acquisition. Finally, the use of a plate-reader in our work limits the ability to use this assay in the field and requires expensive equipment. At the moment, there is no full genomic sequence of *B. zonata*, as well as other pests that might have high sequence similarity in the *Bactrocera* family. The specificity of Cas12a detection relies on genomic data availability, which can contribute to the gRNA design process. This problem also exists with other pests, many of which have no available genomic data or sequence data restricted to the COI gene (Folmer et al. 1994).

Aside from using lateral-flow-based assays or handheld fluorometers as suggested previously, other modifications have been described that can dismiss the use of a plate reader. Moving from the original fluorescence ssDNA reporter (a short ssDNA oligo with a 5’ FAM modification and a 3’ black-hole quencher modification) to more stable and robust reporters such as the coupling of the reporter to gold nanoparticles, gRNA modifications (Fu et al. 2021; Nguyen, Smith, and Jain 2020) or colorimetric reporters (Cao et al. 2021; Y.Zhang et al. 2021). Amplification-free Cas12a methods that utilize complex chemistries to accomplish detection have also been described and may replace the need for RPA or LAMP amplification steps (Choi et al. 2021; Yue et al. 2021). Several insect sequencing efforts are undergoing and hopefully will increase the available genomic data for agriculturally important insect pests. With this increase in sequence availability. We hope our assay will be widely adopted and extended to other pests for one-pot amplification and detection assays in the field and in border controls, enabling rapid, onsite, affordable, and highly specific identification.

## Methods

### Sample collection

All experiments were performed on samples received from the Plant Protection and Inspection Services, Israel.

### gRNA design

gRNAs were designed using CRISPOR (Concordet and Haeussler 2018). Both *B. zonata* and *C. capitata* mitochondrial sequences were obtained from NCBI (accessions NC_027725.1 and NC_000857.1, respectively). Sequences were aligned using the MAFFT version 7 online tool (Katoh, Rozewicki, and Yamada 2019). To find target hotspots within the mitochondrial genome sequences of both insects, we looked for variable regions of up to 500 bp flanked by conserved sequences of at least 30 bp to enable universal RPA amplification. To create unique gRNAs for each insect, both variable regions were concatenated into a single long sequence, and this sequence was used in the CRISPOR gRNA search (Figure 2a). Subsequently, gRNAs were ordered from Integrated DNA Technologies (IDT, gRNA sequences provided in Table S1).

### DNA extraction for gRNA testing

For purification of total DNA from insects, the Qiagen Blood & Tissue kit with the supplementary insect protocol was used (https://www.qiagen.com/kr/resources/resourcedetail?id=cabd47a4-cb5a-4327-b10d-d90b8542421e&lang=en). Specimens were homogenized and then lysed with Proteinase K for 10 min at 56°C. Next, ethanol was added, and the homogenate was loaded onto DNeasy mini columns, washed, and eluted.

### Chelex 100 genomic extraction

Samples were softened in 20 μL ultrapure water (UPW) and ground using a pestle. Next, 100 μL 1×PBS 1% saponin (Sigma-Aldrich, Cat # 8047-15-2) were added, the samples were vortexed, and incubated at RT for 20 min. Lysates were centrifuged for 2 min at 20 G, the supernatant was removed and 100 μL 1 ×PBS was added and centrifuged for 2 min at 20 G. The supernatant was removed and 100 μL 5% Chelex 100 (Bio-Rad, Cat# 1422822) were added. Lysates were vortexed for 5 sec and boiled for 10 min. Next, samples were centrifuged for 1 min at 20 G. The DNA remained in the supernatant, which was subsequently used for RPA amplification.

### RPA amplification

RPA was performed using the TwistAmp^®^ Basic kit (Cat # TABAS03KIT) according to the manufacturer’s protocol. A mix of primers, UPW, MgOAc, and 2 μL from the Chelex reaction were prepared following the protocol concentrations. Next, a lyophilized RPA reaction was suspended with the primer-free rehydration buffer, and the primer mix was added to the reaction, followed immediately by a 20min. incubation at 37°C. 10 μL reactions were similarly performed by keeping the original protocol ratios. Primer sequences can be found in Table S1.

### LbCas12a cleavage assays

All reactions were prepared on ice. gRNA-LbCas12a (NEB, Cat. # M0653T) complexes were prepared by mixing 62.5 nM gRNA with 50 nM LbCas12a in 1XNEBuffer 2.1 to a final volume of 20 μL and incubated in 37°C for 30 min. Next, 1 μM FAM reporter (Supplementary Table 1) and 2 μL of the RPA reaction template were added to the complexes together with 80 μL of 1×NEBuffer 2.1 and incubated for 10 min at 37°C.

Samples were transferred to a black 96 well plate and fluorescence was measured using a Tecan Spark plate-reader with an excitation wavelength of 485 nm and emission was measured at 535 nm. The gain was calibrated to 90.

The detection in the specific primer amplification experiments was performed with a lower concentration of LbCas12a (1μM, NEB, Cat. # M0653S). Interestingly, we noticed a slight improvement in reaction stability using this product.

## Acknowledgments

We thank Dr. Liat Gidron, Dr. Yoav Gazit, and Dr. David Nestel for fly supplies and fruitful discussions. We thank Dr. Karin Mittelman for her insightful comments on the manuscript.

## Contribution

G.P conceived the project, D.M.A, G.P and D.B designed the experiments and wrote the manuscript, and D.M.A and T.P performed the experiments.

## Declarations

No funding was received to assist with the preparation of this manuscript. The authors have no competing interests to declare that are relevant to the content of this article.

---

**Figure S1.**
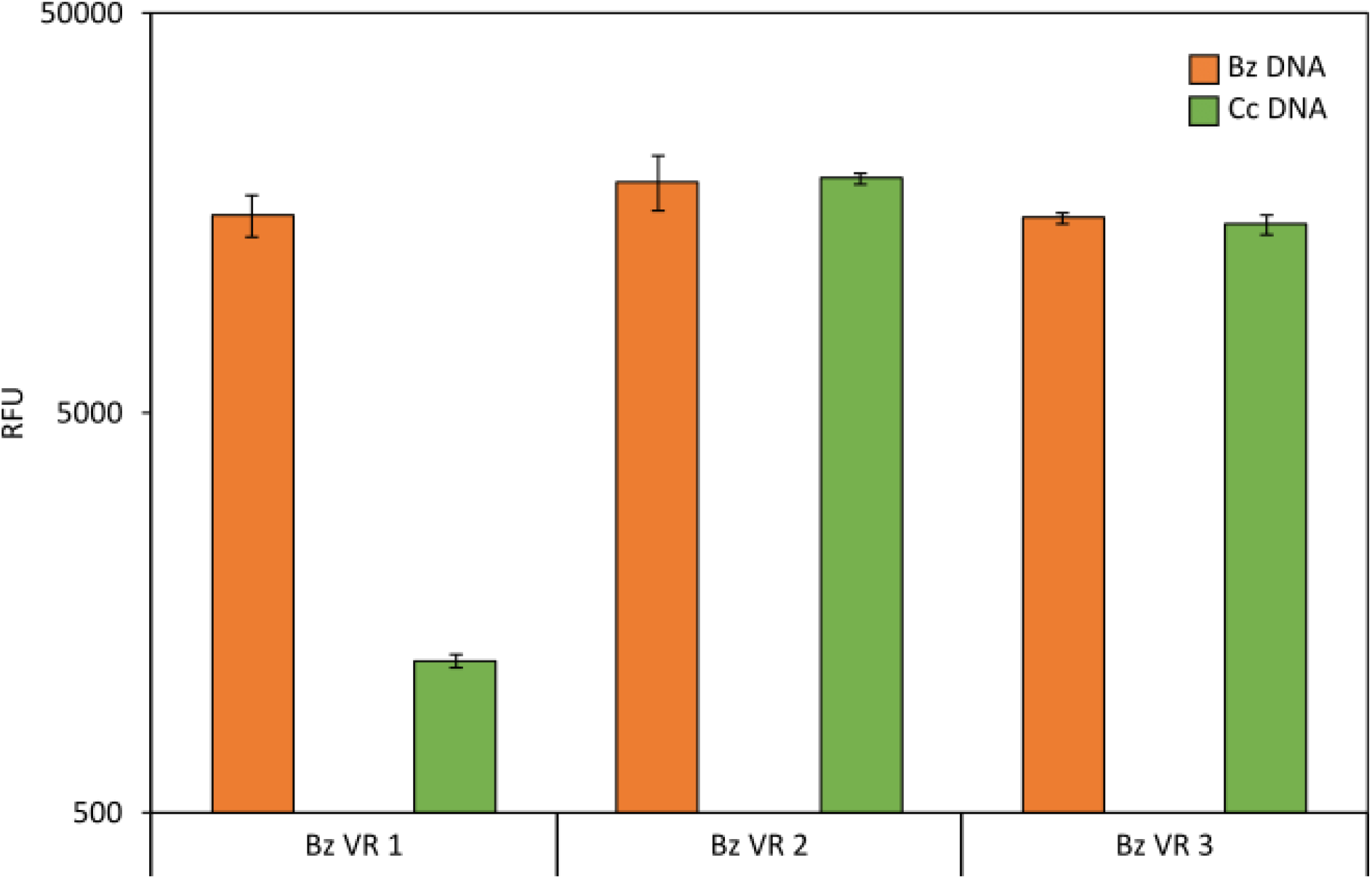
gRNA testing for *B. zonata*. Testing of three different gRNAs based on CRISPOR analysis of the *B. zonata* variable region. *B. zonata* larvae were used, and amplification was performed using PCR with B. *zonata* specific primers (Table S1). Negative control was performed on *C. capitata* larval DNA, amplified with the primers described above. All experiments were performed with three biological repeats and three technical repeats.

**Figure S2.**
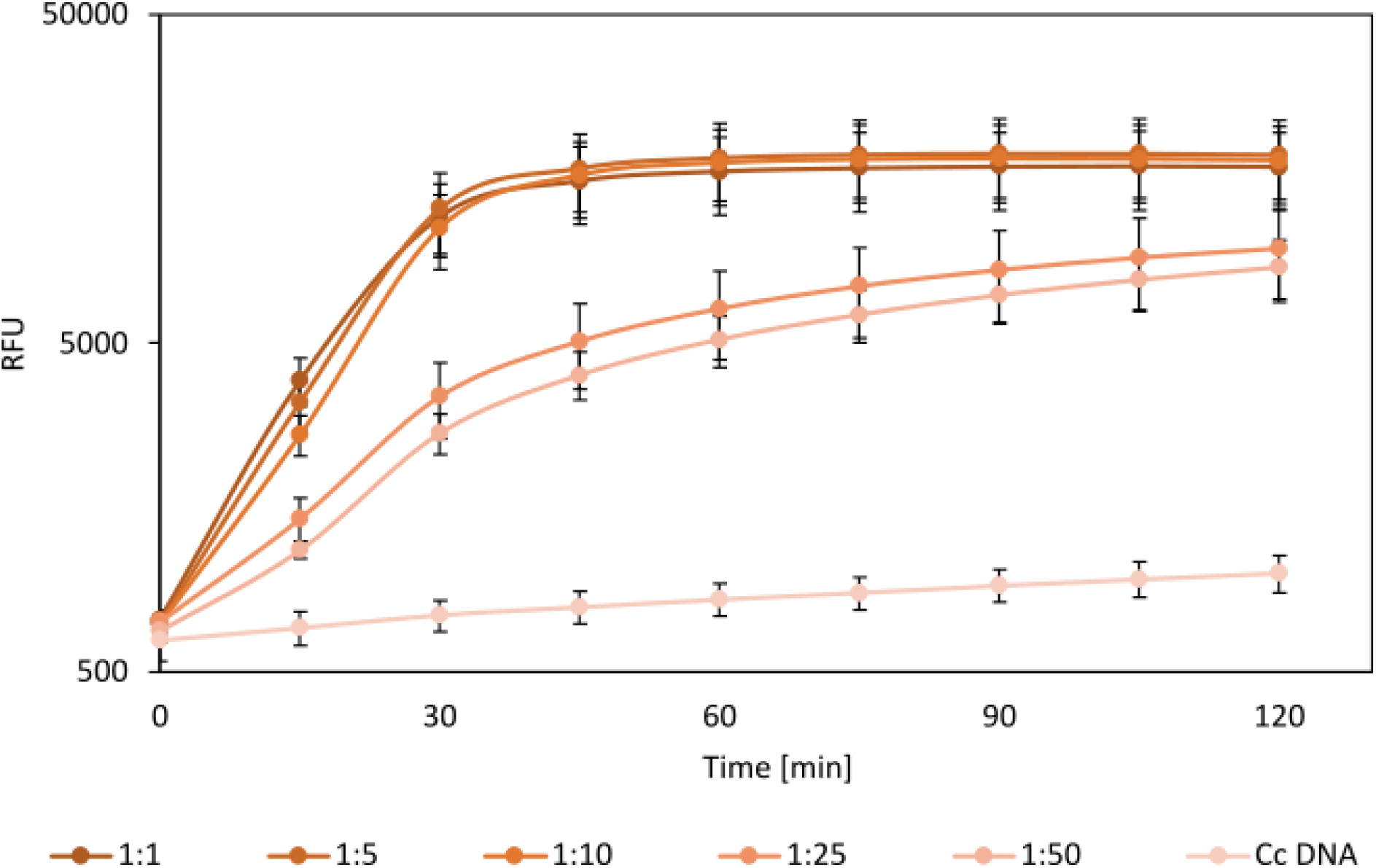
Sensitivity testing for *B. zonata* specific identification. Detection sensitivity of Cas12a-Bz1 gRNA in varying ratios of pooled larvae. Samples containing a single larva of *B. zonata* with increasing amounts of *D. melanogaster* larvae were prepared. DNA was then extracted using Chelex 100, RPA amplified using *B. zonata* specific primers (Table S1), and diagnosed using Cas12a-Bz1 gRNA complexes. As a negative control, *C. capitata* DNA was used. All experiments were performed with three biological repeats and three technical repeats.

**Table S1.**
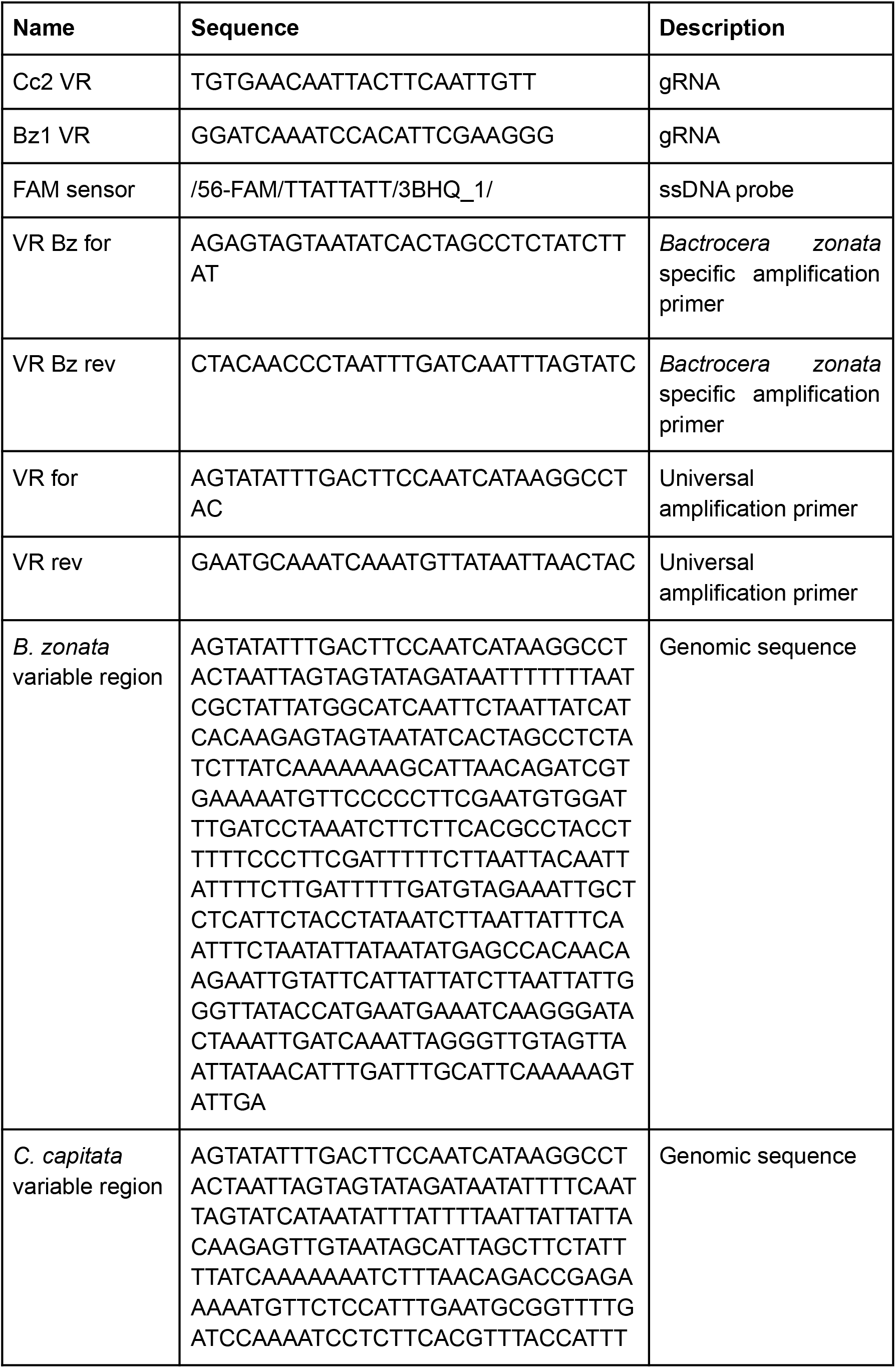
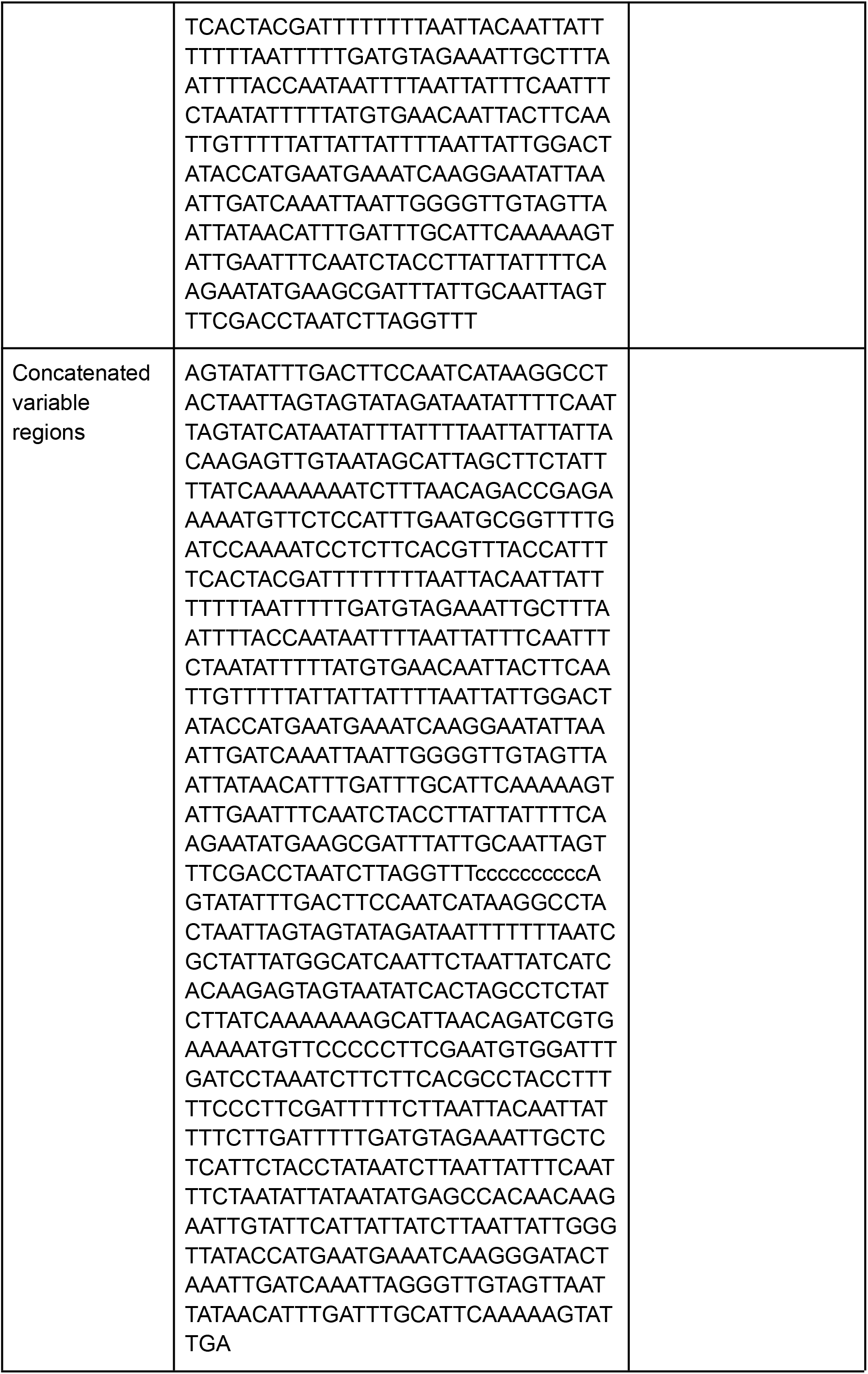
Sequences

**Table S2.**
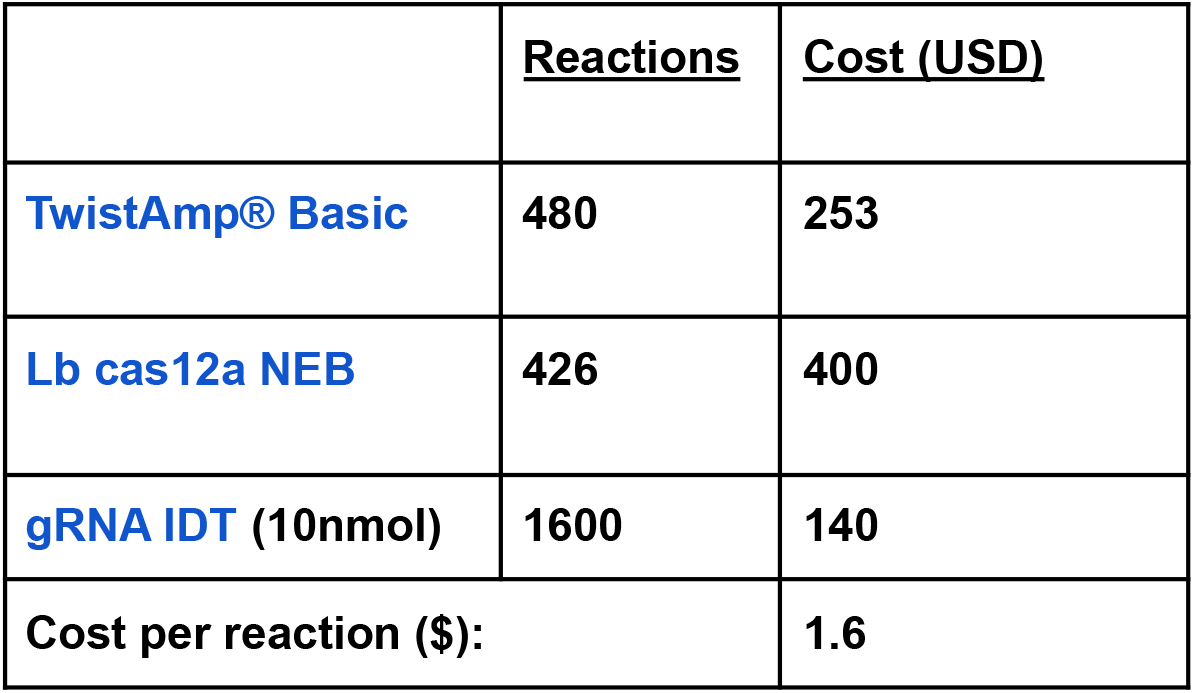
Reagent Costs

