## supplementary figures and tables for "Rapid and sensitive on-site genetic diagnostics of pest fruit flies using CRISPR-Cas12a"

### Supplementary material

#### Figure S1- gRNA testing for *B. zonata*

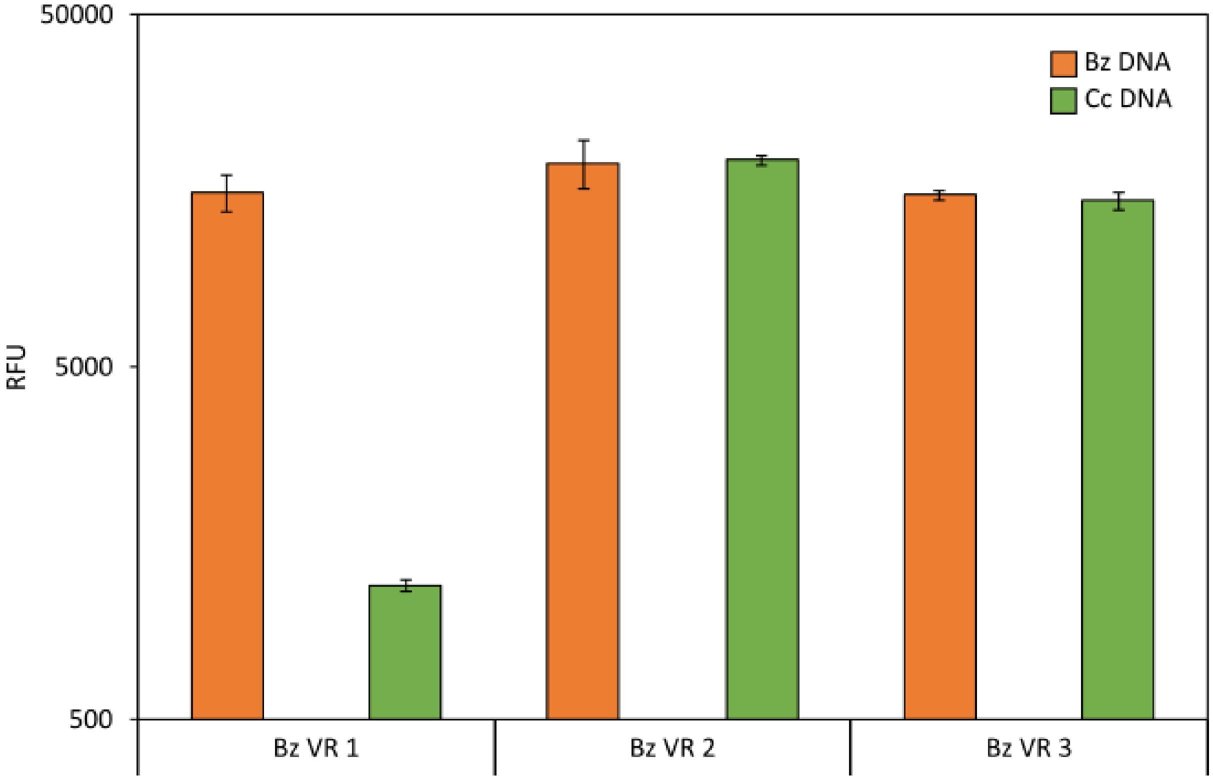

Testing of three different gRNAs based on CRISPOR analysis of the *B. zonata* variable region. *B. zonata* larvae were used, and amplification was performed using PCR with B. *zonata* specific primers (Table S1). Negative control was performed on *C. capitata* larval DNA, amplified with the primers described above. All experiments were performed with three biological repeats and three technical repeats.

#### **Figure S2 - Sensitivity testing for *B. zonata* specific identification**

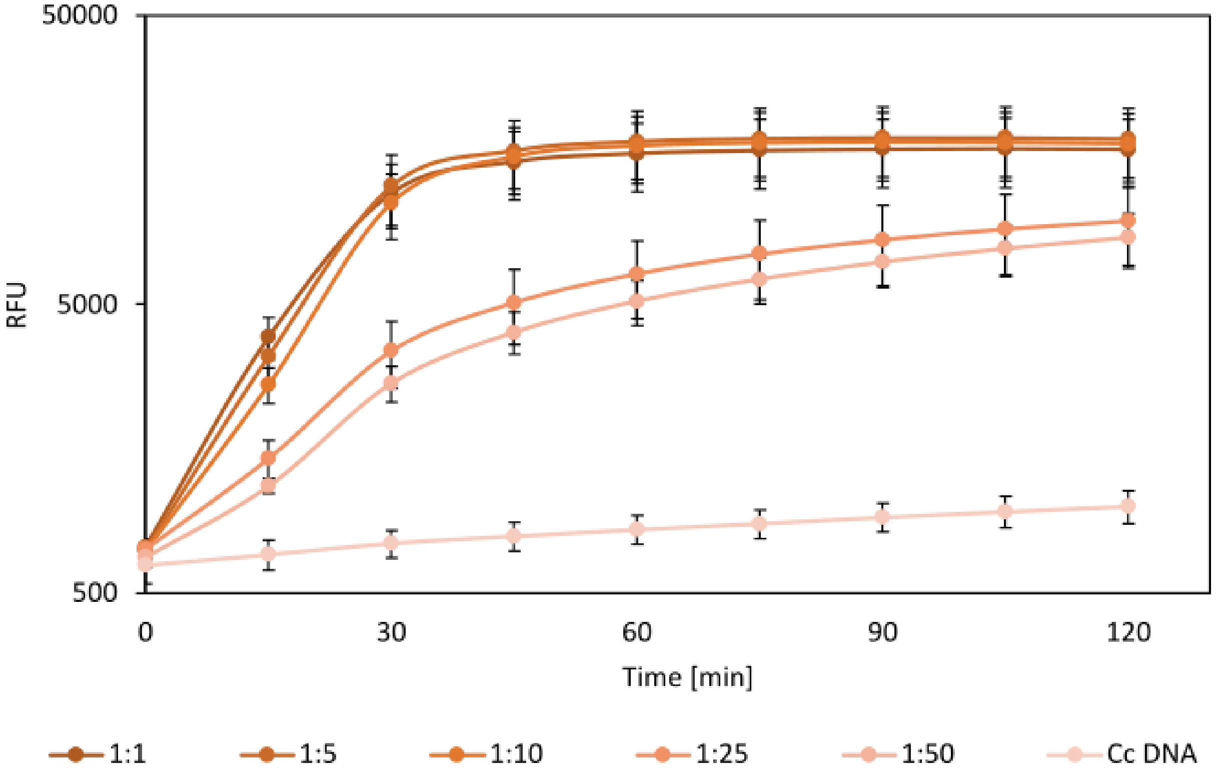

Detection sensitivity of Cas12a-Bz1 gRNA in varying ratios of pooled larvae. Samples containing a single larva of *B. zonata* with increasing amounts of *D. melanogaster* larvae were prepared. DNA was then extracted using Chelex 100, RPA amplified using *B. zonata* specific primers (Table S1) and diagnosed using Cas12a-Bz1 gRNA complexes. As a negative control, *C. capitata* DNA was used. All experiments were performed with three biological repeats and three technical repeats.

Table S1 -DNA Sequences

| Name | Sequence | Description |
|---|---|---|
| Cc2 VR | TGTGAACAATTACTTCAATTGTT | gRNA |
| Bz1 VR | GGATCAAATCCACATTCGAAGGG | gRNA |
| FAM sensor | /56-FAM/TTATTATT/3BHQ_1/ | ssDNA probe |
| VR Bz for | AGAGTAGTAATATCACTAGCCTCTATCTTAT | *Bactrocera zonata* specific amplification primer |
| VR Bz rev | CTACAACCCTAATTTGATCAATTTAGTATC | *Bactrocera zonata* specific amplification primer |
| VR for | AGTATATTTGACTTCCAATCATAAGGCCTAC | Universal amplification primer |
| VR rev | GAATGCAAATCAAATGTTATAATTAACTAC | Universal amplification primer |
| *B. zonata* variable region | AGTATATTTGACTTCCAATCATAAGGCCTACTAATTAGTAGTATAGA TAATTTTTTTAATCGCTATTATGGCATCAATTCTAATTATCATCACAA GAGTAGTAATATCACTAGCCTCTATCTTATCAAAAAAAGCATTAAC AGATCGTGAAAAATGTTCCCCCTTCGAATGTGGATTTGATCCTAAA TCTTCTTCACGCCTACCTTTTTCCCTTCGATTTTTCTTAATTACAATTA TTTTCTTGATTTTTGATGTAGAAATTGCTCTCATTCTACCTATAATCT TAATTATTTCAATTTCTAATATTATAATATGAGCCACAACAAGAATT GTATTCATTATTATCTTAATTATTGGGTTATACCATGAATGAAATCA AGGGATACTAAATTGATCAAATTAGGGTTGTAGTTAATTATAACAT TTGATTTGCATTCAAAAAGTATTGA | Genomic sequence |
| *C. capitata* variable region | AGTATATTTGACTTCCAATCATAAGGCCTACTAATTAGTAGTATAGA TAATATTTTCAATTAGTATCATAATATTTATTTTAATTATTATTACAA GAGTTGTAATAGCATTAGCTTCTATTTTATCAAAAAAATCTTTAACA GACCGAGAAAAATGTTCTCCATTTGAATGCGGTTTTGATCCAAAAT CCTCTTCACGTTTACCATTTTCACTACGATTTTTTTTAATTACAATTAT TTTTTTAATTTTTGATGTAGAAATTGCTTTAATTTTACCAATAATTTT AATTATTTCAATTTCTAATATTTTTATGTGAACAATTACTTCAATTGT TTTTATTATTATTTTAATTATTGGACTATACCATGAATGAAATCAAG GAATATTAAATTGATCAAATTAATTGGGGTTGTAGTTAATTATAAC | Genomic sequence |

| | |
|---|---|
| | ATTTGATTTGCATTCAAAAAGTATTGAATTTCAATCTACCTTATTATT<br>TTCAAGAATATGAAGCGATTTATTGCAATTAGTTTCGACCTAATCTT<br>AGGTTT |
| Concaten-<br>ated<br>variable<br>regions | AGTATATTTGACTTCCAATCATAAGGCCTACTAATTAGTAGTATAGA<br>TAATATTTTCAATTAGTATCATAATATTTATTTTAATTATTATTACAA<br>GAGTTGTAATAGCATTAGCTTCTATTTTATCAAAAAAATCTTTAACA<br>GACCGAGAAAAATGTTCTCCATTTGAATGCGGTTTTGATCCAAAAT<br>CCTCTTCACGTTTACCATTTTCACTACGATTTTTTTTAATTACAATTAT<br>TTTTTTAATTTTTGATGTAGAAATTGCTTTAATTTTACCAATAATTTT<br>AATTATTTCAATTTCTAATATTTTTATGTGAACAATTACTTCAATTGT<br>TTTTATTATTATTTTAATTATTGGACTATACCATGAATGAAATCAAG<br>GAATATTAAATTGATCAAATTAATTGGGGTTGTAGTTAATTATAAC<br>ATTTGATTTGCATTCAAAAAGTATTGAATTTCAATCTACCTTATTATT<br>TTCAAGAATATGAAGCGATTTATTGCAATTAGTTTCGACCTAATCTT<br>AGGTTTccccccccccAGTATATTTGACTTCCAATCATAAGGCCTACTA<br>ATTAGTAGTATAGATAATTTTTTTAATCGCTATTATGGCATCAATTC<br>TAATTATCATCACAAGAGTAGTAATATCACTAGCCTCTATCTTATCA<br>AAAAAAGCATTAACAGATCGTGAAAAATGTTCCCCCTTCGAATGTG<br>GATTTGATCCTAAATCTTCTTCACGCCTACCTTTTTCCCTTCGATTTT<br>TCTTAATTACAATTATTTTCTTGATTTTTGATGTAGAAATTGCTCTCA<br>TTCTACCTATAATCTTAATTATTTCAATTTCTAATATTATAATATGAG<br>CCACAACAAGAATTGTATTCATTATTATCTTAATTATTGGGTTATAC<br>CATGAATGAAATCAAGGGATACTAAATTGATCAAATTAGGGTTGTA<br>GTTAATTATAACATTTGATTTGCATTCAAAAAGTATTGA |

Table S2 - Reagent Costs

| - | **Reactions** | **Cost (USD)** |
|---|---|---|
| TwistAmp® Basic | 480 | 253 |
| Lb cas12a NEB | 426 | 400 |
| gRNA IDT (10nmol) | 1600 | 140 |
| Cost per reaction ($): | | 1.6 |
